# Regulated delivery controls Drosophila Hedgehog, Wingless and Decapentaplegic signaling

**DOI:** 10.1101/2020.08.12.247759

**Authors:** Ryo Hatori, Thomas B. Kornberg

## Abstract

Morphogen signaling proteins disperse across tissues to activate signal transduction in target cells. We investigated dispersion of Hedgehog (Hh), Wingless (Wg), and Bone morphogenic protein homolog Decapentaplegic (Dpp) in the Drosophila wing imaginal disc, and found that delivery to targets is regulated. Cells take up <5% Hh produced, and neither amounts taken up nor extent of signaling changes under conditions of Hh production from 50-200% normal amounts. Similarly, cells take up <25% Wg produced, and variation in Wg production from 50-700% normal has no effect on amounts taken up or signaling. Similar properties were observed for Dpp. Wing disc-produced Hh signals to disc-associated tracheal and myoblast as well as an approximately equal number of disc cells, but the extent of signaling in the disc is unaffected by the presence or absence of the tracheal cells and myoblasts. These findings show that target cells do not take up signaling proteins from a common pool and that both the amount and destination of delivered morphogens are regulated..

**Summary:** The extent of Hh, Wg, and Dpp signaling is independent of the amount of signal produced or the number of recipient cells.

## Introduction

Signaling by morphogen proteins controls many aspects of development, homeostasis and disease (Garcia et al., 2018; Tabata and Takei, 2004; Taipale and Beachy, 2001). These signaling proteins are released from cells that produce them, and they distribute across the tissues they target, forming concentration gradients that induce signal transduction and activate gene expression in a concentration-dependent manner. The importance of regulation by morphogen gradients to growth, cell fate and patterning underlies the imperative to understand how morphogens disperse across tissues.

For more than a century, it has been assumed that morphogens spread across tissues by passive diffusion in extracellular space (either “free” or “restricted”), and both experimental observations and theoretical modeling have been offered in support (Rogers and Schier, 2011). Spreading morphogen proteins have been proposed to exist in various forms, including as multimeric complexes or encapsulated in lipoprotein particles, exosomes, or micelles (Christian, 2012). Implicit in these models are the ideas that signaling is proportional to amounts of signaling proteins produced by designated groups of cells, and that release creates an extracellular pool of signaling protein that distributes in extracellular fluid in ways that are dependent on interactions with substances that are encountered or until they are removed from the pool by degradation or by receptor-mediated absorption. The pool is assumed to be formed by constitutive release from producing cells.

An alternative mechanism of dispersion is direct exchange at cell-cell contacts made by specialized filopodia called cytonemes (Kornberg, 2016). Cytonemes link signaling and target cells with synaptic contacts, and provide conduits that transport signaling proteins between cells. Genetic conditions that impair cytonemes diminish both cytoneme contacts and signaling. Cytoneme synapses have features and attributes that are characteristic of neuronal synapses, including protein composition, close pre- and postsynaptic membrane apposition, voltage sensitivity, and calcium dependence (Huang et al., 2019; Roy et al., 2014). At a neuronal synapse, signaling is titrated by frequency and quantity of neurotransmitter release and on efficiency of neurotransmitter clearance from the synaptic gap (Blakely and Edwards, 2012). It is not known if cytoneme-mediated morphogen signaling at cytoneme synapses is also dependent on regulated release.

Hh, Wg, and Dpp are evolutionarily conserved morphogen signaling proteins that have been implicated in organogenesis and stem cell maintenance, and their mis-regulation in mammals has been linked to inherited diseases and cancers (Briscoe and Thérond, 2013; Morikawa et al., 2016; Nusse and Clevers, 2017). In the columnar cells of the Drosophila wing imaginal disc, Hh is expressed specifically and uniformly by posterior compartment cells (Fig. 1A). In the wing blade primordium of the wing disc, Hh released by posterior compartment cells is taken up by anterior compartment cells within 30 μm (ten cells) of the anterior/posterior (A/P) compartment border. Transfers of Hh from the posterior to anterior compartment cells is cytoneme-dependent (Bischoff et al., 2013; Chen et al., 2017). Hh in the anterior compartment distributes to form a concentration gradient that induces signal transduction and activates expression of target genes in partially overlapping stripes (Callejo et al., 2011; Chen et al., 2017). These domains of expression reflect graded responses to Hh, from highest and “short-range” (*engrailed* (*en*), *patched* (*ptc*), and *dpp*) to lowest and “long-range” (*cubitus interruptus* (*ci*). The spatial relationships of these domains are reproducible, with single cell resolution.

**Figure 1.**
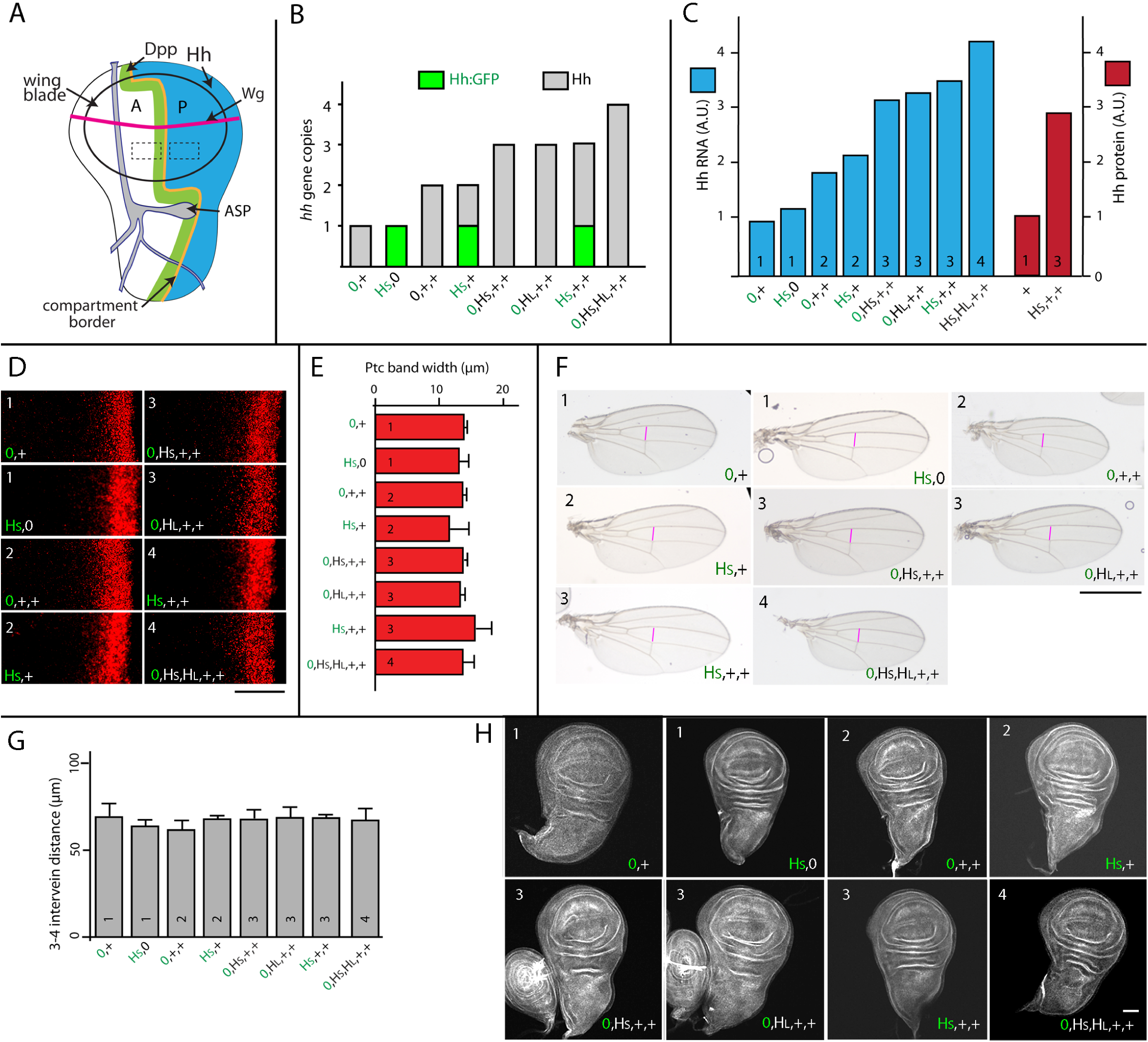
Signal transduction is constant in conditions that vary amounts of Hh production. (A) Schematic of the wing disc and ASP indicating anterior (A) and posterior (P) compartments and showing domains of expression for Hh (blue), Dpp (green), and Wg (magenta). Rectangles (dashed lines, 40 μm x 20 μm) indicate regions that were imaged at high magnification in (D) and in Fig. 2 (A,D). (B) Bar graph showing the number of *hh* genes in genotypes with different combinations of WT *hh* and *hh* BAC transgenes; gray and green bars represent genes encoding Hh and Hh:GFP, respectively. (C) Bar graph showing the amount of Hh RNA (blue) in wing discs and Hh protein (red) in wing disc posterior compartments, measured by qPCR and *α*-Hh antibody staining, respectively, with indicated genotypes and with the number of *hh* genes indicated by numbers in the bars; values normalized to the amount of *hh* RNA and Hh protein in genotype with 1 copy of WT *hh* (0,+). (D) α-Ptc antibody staining in region indicated in (A) by rectangle in A compartment for indicated genotypes. Scale bar, 20μ. (E) Bar graph showing width of the antibody stained Ptc domain in (D). No statistically significant differences (P values>0.05), n = 6-8 for each genotype. (F) Adult wings for indicated genotypes. Scale bar: 100μm. (G) Bar graph showing the measured distance (magenta lines) between 3-4 intervein for adult wings for indicated genotypes. No statistically significant differences (P values>0.05), n = 12-18 for each genotype. (H) Wing discs for each indicated genotype. Error bars in (D,G) indicate Standard Deviation (SD) Scale bar: 100μm. Genotypes: Black letters denote untagged Hh and green letters denote Hh:GFP; H_S_- 40k Hh BAC; H_L_- 100k Hh BAC; +- WT *hh* gene; 0-absence of *hh* gene. 0,+ (*hh*^*AC*^/+); H_S_, 0 (*Hh:GFP 40k BAC*; *hh*^*AC*^/*hh*^*AC*^); 0,+,+ (+ / +); H_S_, + (*Hh:GFP 40k BAC*; *hh*^*AC*^/+); 0,H_S_, +,+ (*Hh 40k BAC*; +/+); H_L_, +,+ (*Hh 100k BAC*; +/+); H_S_, +,+ (*Hh:GFP 40k BAC*; +/+); 0,H_S_,H_L_,+,+ (*Hh 40k BAC* / *Hh 100k BAC* +/+).

In the wing blade primordium, cells that express *dpp* form a stripe of 6-8 cells adjacent to the A/P compartment border (Teleman and Cohen, 2000). *wg* is expressed in a two cell-wide stripe that is orthogonal to the Dpp stripe and straddles the dorsal/ventral (D/V) compartment border (Neumann and Cohen, 1997). Both Dpp and Wg disperse to form concentration gradients on both sides of their respective stripes of expressing cells. Transport of both Dpp and Wg is cytoneme-mediated (Huang and Kornberg, 2015; Roy et al., 2014; Stanganello and Scholpp, 2016).

Here, we asked if distributions of Hh, Wg and Dpp in cells of the wing disc that take up these signaling proteins are proportional to amounts the wing disc produces, and therefore consistent with constitutive release. We also asked if the three target tissues that respond to disc-produced Hh take up Hh from a common pool. Our data show that delivery of signaling proteins to target cells is regulated with respect to both amount and destination.

## Results

### Relationship between Hh production and Hh signaling in the wing disc

Neurotransmitters that are made, packaged, and stored in presynaptic compartments are functionally inert, their precisely controlled release and delivery for juxtacrine activation a signature property of synaptic signaling. In order to investigate whether release of Hh might be regulated at cytoneme synapses, we analyzed Hh signaling in genotypes that express different amounts of Hh. We tested whether amounts of Hh and of Hh signaling in recipient cells are proportional to Hh production, as might be expected of constitutive, unregulated release by producing cells, or if they are independent of production as might be expected of regulated release and delivery.

We first monitored *hh* RNA in wing discs with genotypes that vary the number of wildtype (WT) *hh* genes and *hh* transgenes (Fig. 1B). The *hh* transgenes are BAC plasmids containing either the WT *hh* transcription unit in a genomic fragment of 40kb (HS) or 101 kb (HL), or HS, a 40kb genomic fragment into which GFP has been recombined in frame (Hh:GFP) (Chen et al., 2017). Flies without a functional *hh* gene die as embryos, but haploid flies with only one BAC transgene (encoding Hh (HS, HL, or HS) have normal appearance and wing discs have normal morphology (Chen et al., 2017). Hh:GFP encoded by this transgene is therefore presumed to be a functional surrogate for the normal, WT protein. We used qPCR to measure amounts of *hh* RNA in animals with 1, 2, 3, or 4 *hh* genes, and determined that the amount of *hh* RNA in wing discs scales with gene dosage (Fig. 1C). These results are consistent with the idea that *hh* RNA expression is directly proportional to gene copy number.

To investigate how production and distribution of Hh protein scale in the wing disc and how Hh production correlates with signaling, we measured Hh amounts in the wing blade primordium by monitoring Hh immunohistochemically with *α*-Hh antibody. In discs with either one or three *hh* genes, the amount of Hh detected in the Hh-producing cells increases 2.9 times (Fig. 1C). This result is consistent with the qPCR analysis, and with the idea that both Hh RNA and protein scale with gene dosage.

To determine if different amounts of Hh expression alter growth and patterning, we examined several parameters that respond to and are sensitive to Hh signaling: expression of Hh gene targets, size and shape of the wing and wing disc, and wing vein pattern. Expression of the *ptc* gene in the wing disc is up-regulated by Hh signaling in a band of cells at the A/P compartment border. The width of this band decreases under conditions in which Hh signal transduction is reduced specifically in the responding cells (Molnar et al., 2011), and increases under conditions in which Hh signaling is elevated (Cheng et al., 2012; Wang and Holmgren, 1999). In discs with 1, 2, 3, or 4 *hh* gene copies, we did not detect differences in the size of the Ptc band (Fig. 1 D,E). In the adult wing, the size of the intervein region between veins 3 and 4 is sensitive to Hh signaling, decreasing under conditions of low Hh signaling and increasing under conditions of elevated levels (Casso et al., 2011; Mullor et al., 1997; Strigini and Cohen, 1997). We did not detect changes in the size or shape of either the wing disc, adult wing or 3-4 intervein in flies with 1, 2, 3, or 4 *hh* gene copies (Fig. 1F,G).

To characterize the apparent insensitivity of the disc and wing to different amounts of Hh production, we consider two possibilities. If the amount of Hh taken up by the recipient, target cells is proportional to production, each recipient cell might scale the outputs of Hh signal transduction relative to its neighbors. This mechanism might adjust relative responses independently of absolute amounts, determining growth and pattern by the slope of the concentration gradient across the field of cells. This type of mechanism was proposed for the morphogen gradient of Dpp in order to model the effects of mosaic ectopic activation induced by expression of a constitutively active Dpp receptor (Rogulja and Irvine, 2005). Alternatively, the amount of Hh released from producing cells might be regulated so that a recipient cell receives an amount of Hh that is not dependent on the amount produced. To distinguish between these mechanisms, we quantified Hh taken up by recipient cells in the wing blade primordium.

We first monitored Hh histologically with *α*-Hh antibody in genotypes with 1 (1xWT), 2 (2xWT), 3 (2xWT, 1xBAC), or 4 (2xWT, 2xBAC) *hh* genes. Images of small rectangular regions of the anterior and posterior compartments with these genotypes (Fig. 1A) show that Hh amounts increase proportionally with gene dosage in the posterior compartment where Hh is produced. This result is consistent with the qPCR and histological analyses shown in Figure 1C, and with the idea that both Hh RNA and protein scale with gene dosage. Images of the same small rectangular region in the anterior compartment with these genotypes show that Hh amounts in anterior compartments are not detectably different with 1, 2, 3, or 4 *hh* genes (Fig. 2A,B)., but that Hh amounts in anterior compartments are not detectably different with 1, 2, 3, or 4 *hh* genes (Fig. 2A,B). Quantification of Hh detected by antibody staining in the entire wing primordium of normal (2xWT) discs shows that approximately 5.2% is in the anterior compartment (n=7; standard deviation=1.2%). These results are consistent with the idea that most Hh produced in the posterior compartment is not released (and does not signal), and that Hh export is not linked directly to production.

**Figure 2.**
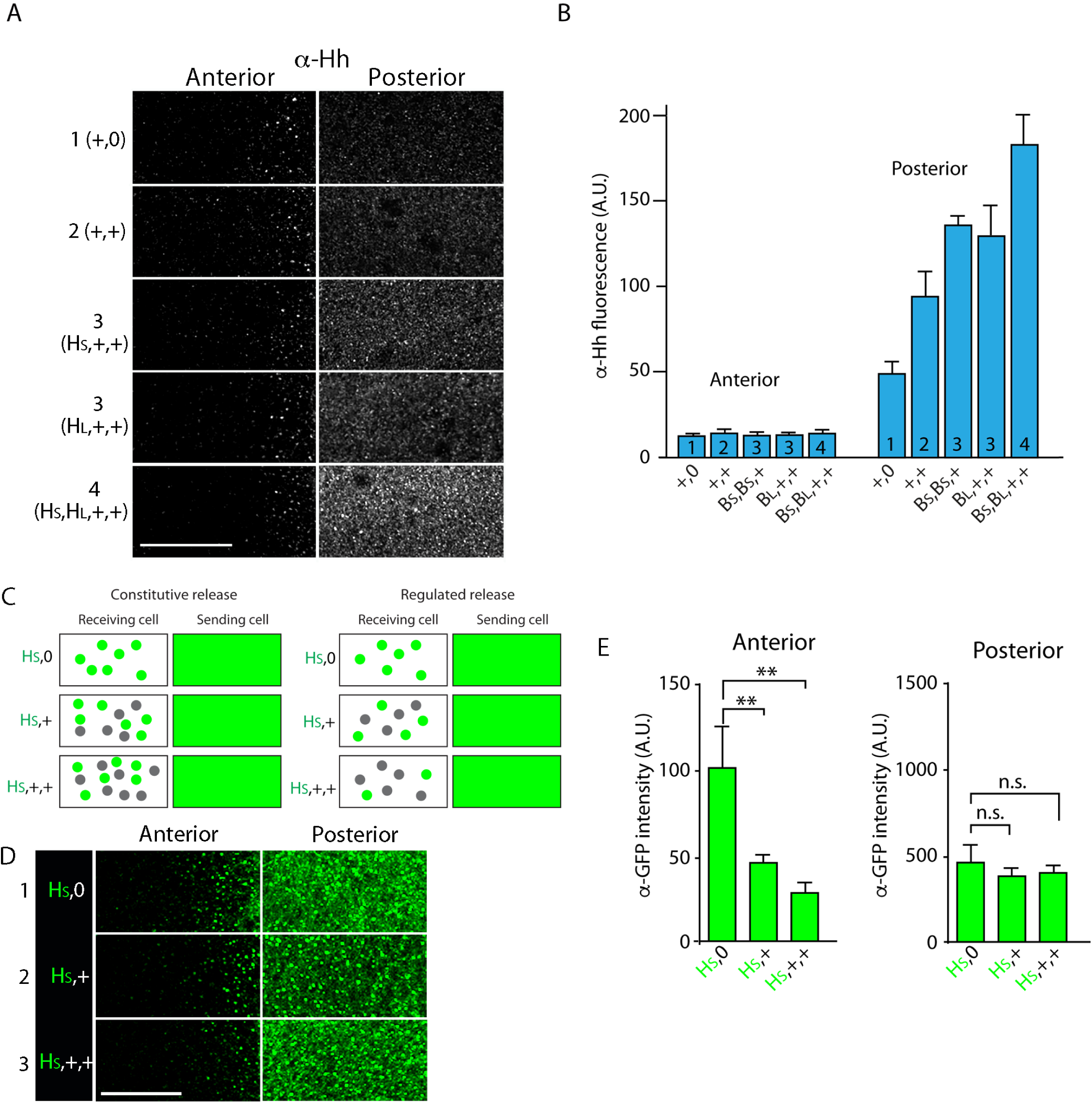
Hh delivery is constant in conditions that vary amounts of Hh production. (A) α-Hh antibody staining in regions indicated in Fig. 1A for indicated genotypes. (B) Bar graph showing the intensity of α-Hh antibody staining in A and P compartments of wing blades for indicated genotypes. No statistically significant differences for A compartment (P values>0.05); for P compartment, staining was statistically different for genotypes with different numbers of *hh* genes (1, 2, 3, or 4; student’s t-test, P<0.05), but not between equivalent numbers (3 and 3; P>0.05), n = 5-7 discs for each genotype. (C) Schematic portraying the predicted differences between constitutive release and regulated release for different genotypes, Hh and Hh:GFP indicated by gray and green dots, respectively. (D) Images of α-GFP antibody staining in regions indicated in Fig. 1A for indicated genotypes. (E) Bar graphs showing α-GFP antibody staining for indicated genotypes. **-P value<0.005, n.s.-P value>0.05; n = 4-6 for each genotype. Abbreviations as in Fig. 1. Scale bars: 20 μm.

To characterize the relationship between Hh production and delivery further, we used *α*-GFP antibody to analyze genotypes with one Hh:GFP-encoding BAC transgene (HS) together with either zero, one, or two (untagged) WT Hh genes (Total genes: 1: HS; -/-; 2: HS; -/+; and 3: HS; +/+) (Fig. 1B). As depicted in the Figure 2C drawing, *α*-GFP antibody staining of GFP-tagged Hh that is titrated with different amounts of untagged Hh distinguishes between constitutive and regulated delivery in these genotypes. If delivery of Hh to the anterior compartment is proportional to gene dosage and not regulated, Hh:GFP amounts in the anterior compartment are expected to be unaffected by co-production of untagged Hh so that the Hh:GFP remains constant as gene dosage and production increases. However, if delivery is regulated, the fraction of Hh:GFP in the anterior compartment is expected to decrease as the fraction of untagged Hh increases in proportion to total gene copy.

Analysis of wing discs with one Hh:GFP BAC (HS) and 0, 1, or 2 WT *hh* genes shows that Hh:GFP in the producing cells of the posterior compartment is not diminished by the presence of *hh* genes that encode untagged Hh (Fig. 2D,E). This is consistent with the idea that production of both Hh RNA and protein are proportional to gene copy. In contrast, Hh:GFP amounts in the Hh-receiving cells of the anterior compartment decreases in proportion to number of *hh* genes that encode untagged Hh. This shows that Hh:GFP was diluted by the presence of untagged Hh. This result is consistent with the amounts of Hh we detected in the anterior compartment with *α*-Hh antibody in genotypes with 1, 2, 3, or 4 *hh* genes (Fig. 2A,B). We conclude that the amount of Hh delivered to the anterior compartment is constant and does not scale with production.

### Dpp production and signaling in the wing disc

To investigate whether regulated export is also a feature of Dpp signaling, we monitored signaling and dispersion of Dpp in genotypes with different numbers of *dpp* genes. We created a Dpp-encoding BAC transgene (BD) that rescues *dpp* haploinsufficiency: animals with one WT *dpp* and one BD (*+/dpp*^*H46*^; *+/*BD) are viable and their wing size is comparable to WT flies (Fig. 3A,B), indicating that the Dpp BAC is a functional substitute for a WT *dpp* gene. To monitor different amounts of *dpp* expression, we compared wing discs with two or four copies of *dpp* gene (2 copies: +/+; 4 copies: +/+; *BD/BD*). First, to examine proportionality between *dpp* gene copy and Dpp protein, we stained wing discs with antibody that recognizes the prodomain of unprocessed Dpp (Akiyama and Gibson, 2015; Panganiban et al., 1990). Staining in cells that produce Dpp was approximately double in the four copy compared to the two copy genotype (Fig. 3C,D). Second, we examined wing size, which is sensitive to different amounts of Dpp signaling. Mutant conditions that decrease Dpp signal transduction reduce wing disc growth and mutant conditions that elevate signal transduction cause overgrowth (Capdevila and Guerrero, 1994; Spencer et al., 1982). We found that wing size did not differ between genotypes with two or four *dpp* genes (Fig. 3A,B). Third, we asked if signal transduction increases with gene dosage and Dpp production. *α*-pMAD staining, a readout of Dpp signaling, forms a band that straddles and flanks *dpp* expressing cells in WT discs. The width of the pMAD-staining stripe was not significantly changed in wing discs with two or four gene copies (Fig. 3E,F), indicating that increased Dpp production does not increase signal transduction. In sum, these results show that the wing disc is insensitive to increased levels of Dpp production.

**Figure 3.**
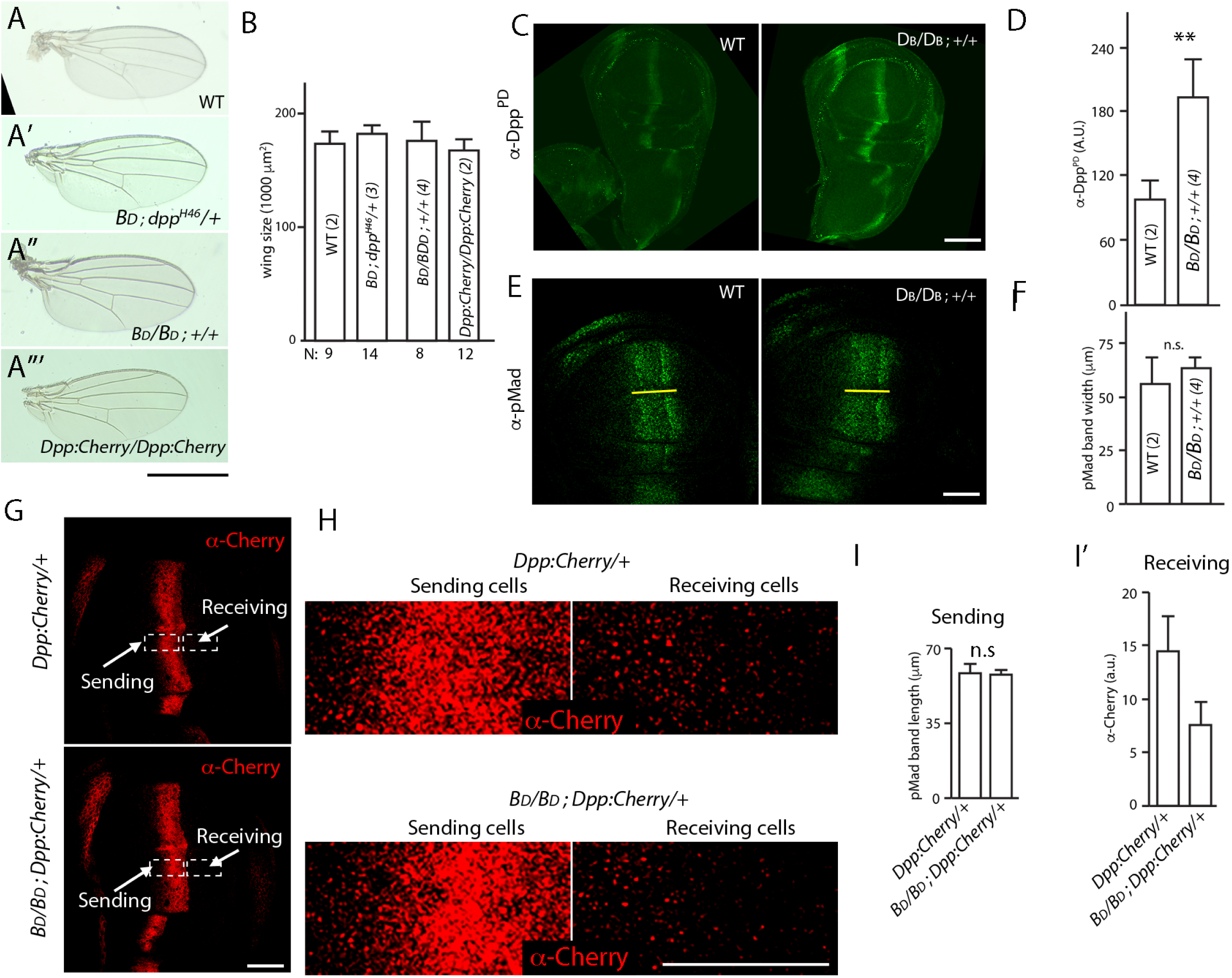
Dpp delivery and signal transduction are constant in conditions that vary amounts of Dpp production. (A-A”‘) Adult wings for indicated genotypes. Scale bar: 100μm. ((B) Bar graph showing size of adult wings for genotypes in (A-A”‘); error bars indicate SD, N indicates number of wings analyzed; no statistically significant differences indicated by student’s t-test (P values>0.05). (C) Wing discs with two (WT) and four *dpp* genes: WT (+/+); four (BD/BD; *+/+*) stained with α-Dpp prodomain (α-DppPD) antibody; scale bar: 100μm. (D) Bar graph quantifying α-DppPD antibody staining for wing discs with indicated genotypes, n = 7 (2 genes) and 8 (4 genes). Difference is statistically significant (student’s t-test P value<0.005). (E) Images of wing discs with indicated genotypes stained with α-pMAD antibody; yellow line marks the width of pMAD band; scale bar: 50μm. (F) Bar graph quantifying α-pMAD antibody staining for wing discs with indicated genotypes; n = 7 (two genes) and 3 (four genes). No statistically significant differences indicated by student’s t-test (P values>0.05) (G) Wing blades with (upper panel) one untagged Dpp (+) and one Dpp:Cherry encoding gene, or (lower panel) three untagged Dpp and one Dpp:Cherry encoding gene stained with α-Cherry antibody; scale bar: 50μm. (H) High magnification image of boxed regions in (J); scale bar: 25μm. (I) Bar graph quantifying α-Cherry antibody staining in sending and receiving regions for indicated genotypes; error bars indicate SD, no statistically significant differences indicated by student’s t-test (P>0.05); n = 7 for each genotype. (M) Same as (J) for receiving cells. Difference is statistically significant (P<0.05). Abbreviations: BD (Dpp-encoding BAC transgene), Dpp:Cherry (Dpp:Cherry knock-in allele).

To monitor Dpp distributions, we examined discs stained with *α*-Dpp antibody to compare genotypes with one Dpp:Cherry knock-in allele and either one or three *dpp* genes that encode untagged protein (Fig. 3G). The experimental setup and rationale are similar to the analysis of Hh:GFP depicted in Figure 2C. Evaluation of the two genotypes showed that Dpp:Cherry amounts in producing cells does not change (Fig. 3H,I), and that amounts of Dpp:Cherry in non-producing, receiving cells decreased in proportion to the number of genes that encode untagged Dpp (Fig. 3I’). This finding, that the amount of Dpp:Cherry exported to target cells decreases as the ratio of tagged:untagged Dpp declines, is consistent with the idea that transmission of Dpp to targets is regulated.

### Wingless production and signaling in the wing disc

We investigated the nature of Wg export by monitoring expression of Wg and the Wg gene target Senseless (Sens) in three genotypes that have different numbers of functional *wg* genes: 1 (*wg*^*+*^*/wg*^-^), 2 (*wg*^*+*^*/wg*^*+*^), and 2^+over-expression^ (*wg-Gal4 UAS-Wg:GFP*; *wg*^*+*^*/wg*^*+*^). α-Wg antibody staining shows that Wg production is proportional to gene copy number in the discs with 1 and 2 *wg* genes, and that the *wg-Gal4* driver generates approximately seven times more Wg than a single endogenous gene Fig. 4A,A’,B). Despite the differences in expression between these genotypes, the amount of Wg in the neighboring cells that receive Wg was unchanged (Fig. 4A,A’,B). The fraction of Wg present in the neighboring cells relative to the total produced in the wing blade therefore decreased with increasing functional gene dosage, from approximately 41% (1 copy) to 24% (2 copies), and 3.5% (7 functional equivalents). *α*-Sens antibody detects two narrow, parallel stripes of expression that are immediately adjacent to but do not overlap the Wg-expressing cells (Fig. 4A”), and the patterns of Sens expression were not detectably different in these genotypes (Fig. 4C,C’). We conclude that Wg transmission is regulated.

**Figure 4.**
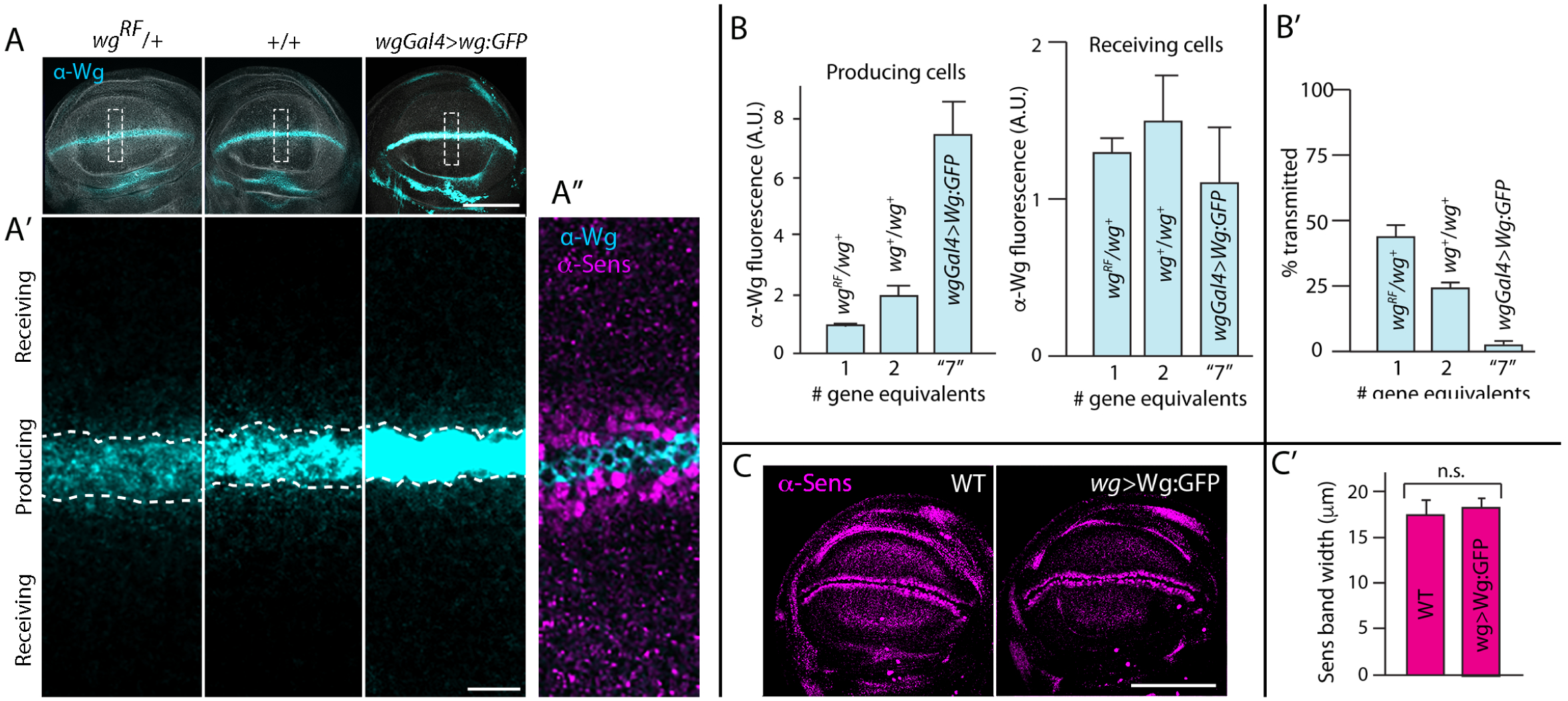
Wg signal transduction is constant in conditions that vary amounts of Wg production. (A-A’) Wing blades for indicated genotypes stained with α-Wg antibody (cyan) and phalloidin (grey); one gene (*wg*^*-*^/+), two genes (+/+), over-expression (*wgGal4>UAS-wg*; *+/+*). Scale bar: 100μm. (A’) higher magnification images of boxed regions (30 μm x 90 μm) in (A), dashed white lines mark boundary between producing and receiving cells. Scale bar: 10μm. (A’’) Optical section of region similar to (A’) stained with α-Wg (cyan) and α-Sens antibodies (magenta). (B) Bar graphs quantifying α-Wg staining for indicated genotypes; n = 5-6 for each genotype. Values are normalized to the intensity of α-Wg staining for *wg*^*RF*^/*wg*^*+*^ (1 copy of wg gene). # gene equivalents indicate approximate Wg production functionality for each genotype. Difference in the producing cells are statistically significant (student’s t-test P value <0.005), while difference in the receiving cells are not (student’s t-test P value >0.05). (B’) Bar graph quantifying fraction of Wg in the receiving cell as % of total wing blade α-Wg antibody intensity in receiving cell. Statistical significance indicated by P<0.0005. (C) Wing blades with two WT genes or Wg over-expression (*wgGal4>UAS-Wg:GFP*; *+/+*) stained with α-Sens antibody. Scale bar: 100μm. (C’) Bar graph quantifies width of α-Sens antibody stained band in maximum intensity projections of optical sections for entire apical-basal depth; ten length measures were taken for each disc; no statistically significant differences (P>0.05); n = 4 for each genotype. Genotypes: wg^-^/+ (*wg*^*RF*^/+); +/+ (WT); *wg-Gal4*>*wg* (*wg-Gal4*; *UAS-Wg:GFP / +/+*).

### Relationship between cytonemes and Hh production

Previous studies in several different systems show that the numbers of cytonemes that cells extend correlate with amount of signal transduction activity. Whereas cells with few cytonemes have low signaling activity, cells with more than normal numbers of cytonemes have elevated levels (Bischoff et al., 2013; Chen et al., 2017, 2017; Du et al., 2018; Huang and Kornberg, 2016; Huang et al., 2019; Mattes et al., 2018; Roy et al., 2011, 2014). To investigate the relationship between cytonemes and Hh production, we monitored ASP cytonemes in genotypes with different numbers of *hh* genes.

We first asked if delivery of Hh to ASP cells is sensitive to amounts of Hh production by monitoring two conditions that are dependent on Hh signaling in the ASP: tissue morphology and expression of *engrailed* (*en*), which is a transcriptional target that is induced by Hh signaling in the wing blade and ASP (Guillén et al., 1995; Hatori, R. and Kornberg, 2020). In WT, the ASP has a proximal narrow stalk and distal bulb (Fig. 1A), and *en* expression is graded, with highest levels in the tip cells (Fig. 5A). In mutant conditions with elevated Hh signaling (e.g. ectopic over-expression of Hh in the ASP), the stalk is absent and En expression extends to more proximal tracheal cells (i.e. the transverse connective; Fig. 1A), whereas in mutant conditions with reduced Hh signaling (e.g. *smoothened* loss-of-function and Patched over-expression), the stalk is elongated and En expression is reduced (Fig. 5B). We found that in genotypes with 1-4 *hh* genes, neither ASP morphology nor extent of En expression changed (Fig. 5A,C). These results show that the ASP is insensitive to different amounts of Hh production and are consistent with the idea that Hh delivery is regulated.

**Figure 5.**
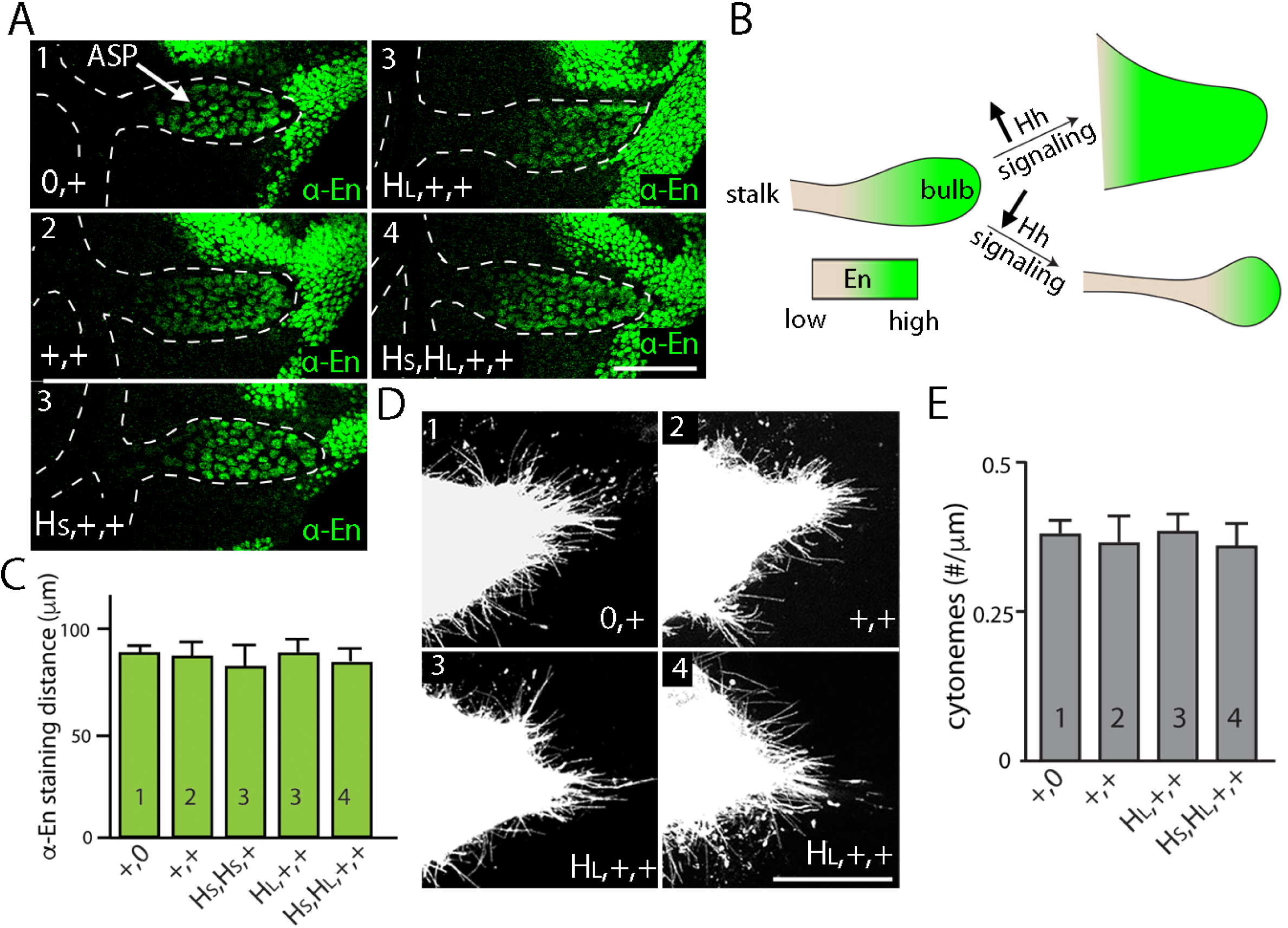
Neither signal transduction nor cytoneme number scales with Hh production in the ASP. (A) Schematic showing En expression (green) in WT ASP (left), ASP with no stalk and high levels of Hh signal transduction (top right), or ASP with elongated stalk and low levels of Hh signal transduction (bottom right). (B) α-En staining (green) of ASPs (bulb within white dashed line) for indicated genotypes (number of *hh* genes indicated in upper left). (C) Bar graph quantifying the distance of α-En antibody staining from the tip of the ASP toward the stalk for indicated genotypes (numbers of genes indicated in bars); no statistically significant differences indicated by P>0.05); n = 5-7 for each genotype. (D) Cytonemes marked by the expression of *Cherry:CAAX* (*btl-lexA>lexO-Cherry:CAAX*) in the ASP for indicated genotypes (number of genes indicated in upper left). (E) Number of cytonemes for indicated genotypes (number of genes indicated in bars). Statistically significant differences indicated by P >0.05; n = 5 for each genotype. Abbreviations as in Fig. 1.

We next investigated the relationship between Hh production and cytonemes. We analyzed the number of cytonemes in genotypes with 1, 2, 3, and 4 *hh* genes by marking ASP cytonemes with membrane-tethered Cherry (*btl>*CD8:Cherry). Cytonemes that extend from the distal tip of the ASP take up Hh and contain Ptc (Chen et al., 2017), and in the experimental genotypes, the number of distal tip cytonemes was not statistically different (Fig. 5D,E). We conclude that the number of cytonemes and amount of cytoneme-mediated Hh uptake are insensitive to conditions that reduce or increase Hh production by a factor of two.

### Expression of modulators of morphogen protein signaling

In addition to the synthesis of signaling proteins by producing cells and pathways of signal transduction in receiving cells, morphogen signaling involves post-translational processes that prepare Hh, Dpp, and Wg in producing cells, feedback regulation in receiving cells, and extracellular proteins that influence activity. We investigated whether changes in the production of Hh, Dpp, or Wg affects the expression of genes that encode functions known to modulate signaling because the expression of these genes might provide feedback regulation that compensates for changes in the amounts of proteins that are released or taken up. We might expect, for example, that the expression of a gene that provides negative feedback increases under conditions of increased signaling protein production. *Shifted* (*Shf*) encodes an extracellular factor that is required for the normal distribution of Hh, and Shf protein levels decrease in conditions of lowered signaling (Glise et al., 2005). We quantified *shf* transcripts in discs with one and four *hh* genes by qPCR, but detected no change in *shf* expression (Fig. 6A,B). This insensitivity to Hh amounts suggests that Shf does not control Hh release. *brinker* (*brk*), *Pentagone* (*Pent*), *Short gastrulation* (*Sog*), and *Crossveinless-2* (*Cv-2*) negatively affect Dpp signaling. Brk is a transcriptional repressor of Dpp signal transduction whose expression is suppressed by Dpp. *Pent, Sog*, and *Cv-2* encode extracellular proteins that bind Dpp and negatively affect spread and signaling (Raftery and Umulis, 2012) (Fig. 6A). Ectopic Dpp signaling suppresses *brk, Pent* and *Sog* and upregulates *Cv-2* expression (Raftery and Umulis, 2012; Yu et al., 1996). qPCR analysis detected no changes to expression of *brk, Pent, Sog*, or *Cv-2* in genotypes with 2 or 4 *dpp* genes (Fig. 6B). *Notum* expression is induced by Wg signaling and encodes an extracellular deacylase of Wg that inhibits Wg signaling (Minami et al., 1999), but its expression is not influenced by changes to Wg gene number (Fig. 6B). In sum, these data do not support the idea that expression of known modulators of the Hh, Dpp, and Wg pathways compensate for changes in amounts of morphogen production and are consistent with the idea that release is regulated.

**Figure 6.**
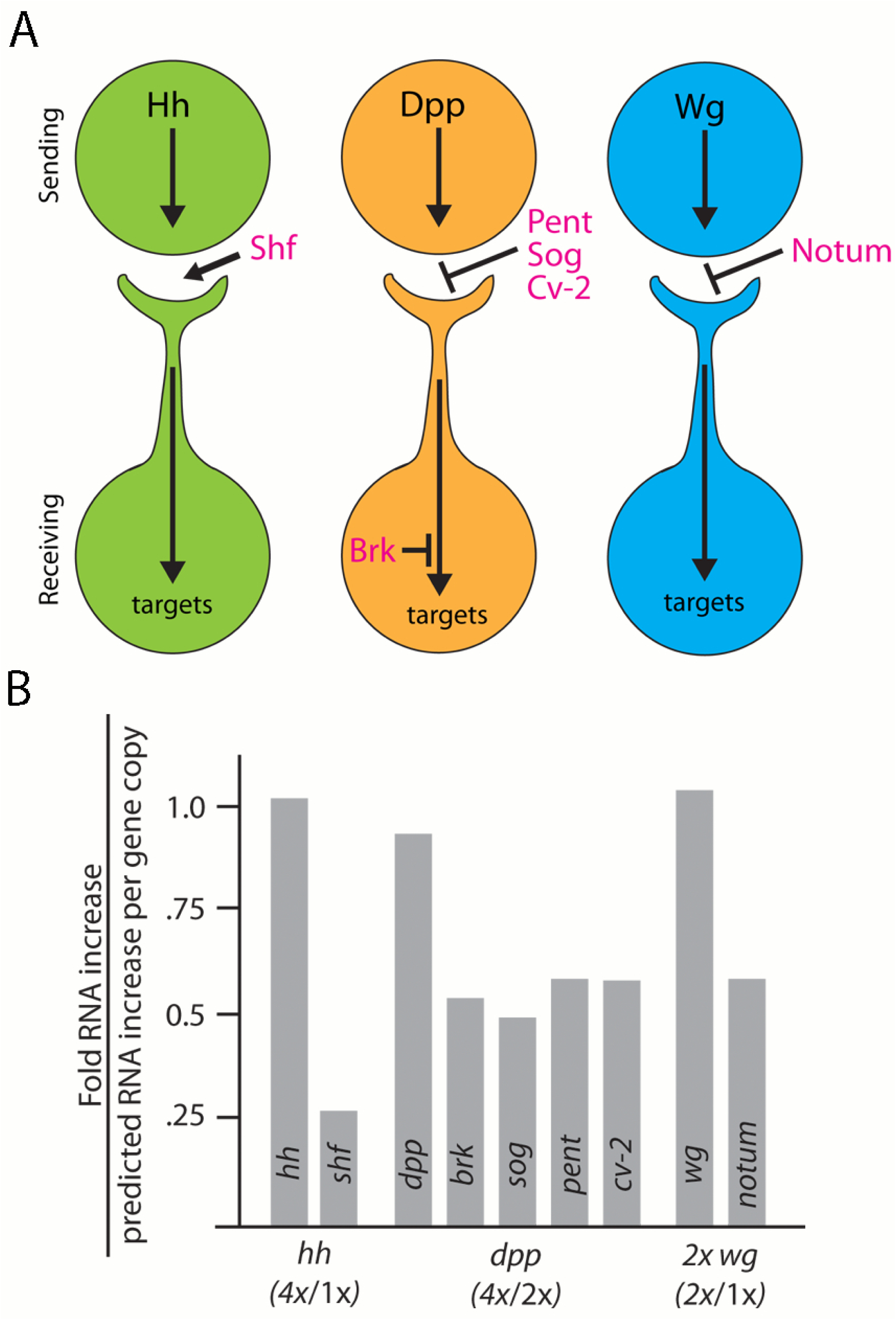
Expression of morphogen signaling modulators is not affected by varying amounts of morphogen production. (A) Schematic showing where morphogen signaling modulators are predicted to function in the context of cytoneme-mediated exchange. Shf, an extracellular factor that facilitates Hh dispersion; Pent, Sog, and Cv-2, extracellular inhibitors of Dpp signaling; Brk, a transcriptional repressor of Dpp signal transduction; Notum, an extracellular inhibitor of Wg signaling. (B) Bar graph showing the levels of morphogen signaling modulator mRNA as determined by qPCR. Bars represent ratio between change in mRNA levels relative to predicted RNA increase that scales with gene copy.

### Hh gradients form independently in the wing disc, ASP, and myoblasts

To characterize how signaling proteins are apportioned among the cells they target, we analyzed Hh signaling in a uniquely positioned group of Hh-responding cells near the Hh-producing cells of the wing disc notum primordium. Cells in this region include cells of the wing disc anterior compartment, the ASP and myoblasts, and because of the close proximity of these cells to Hh-producing cells in the disc, and because no other Hh-producing cells are as close, we presume that the Hh they receive originates from this one source (Hatori, R. and Kornberg, 2020). We designated an area 150 μm x 150 μm that includes all the Hh-responding cells in the notum, ASP, and myoblasts, as a Hh “microenvironment” (Fig. 7A-C). We monitored Hh signaling in this region by Ptc expression and determined that

**Figure 7.**
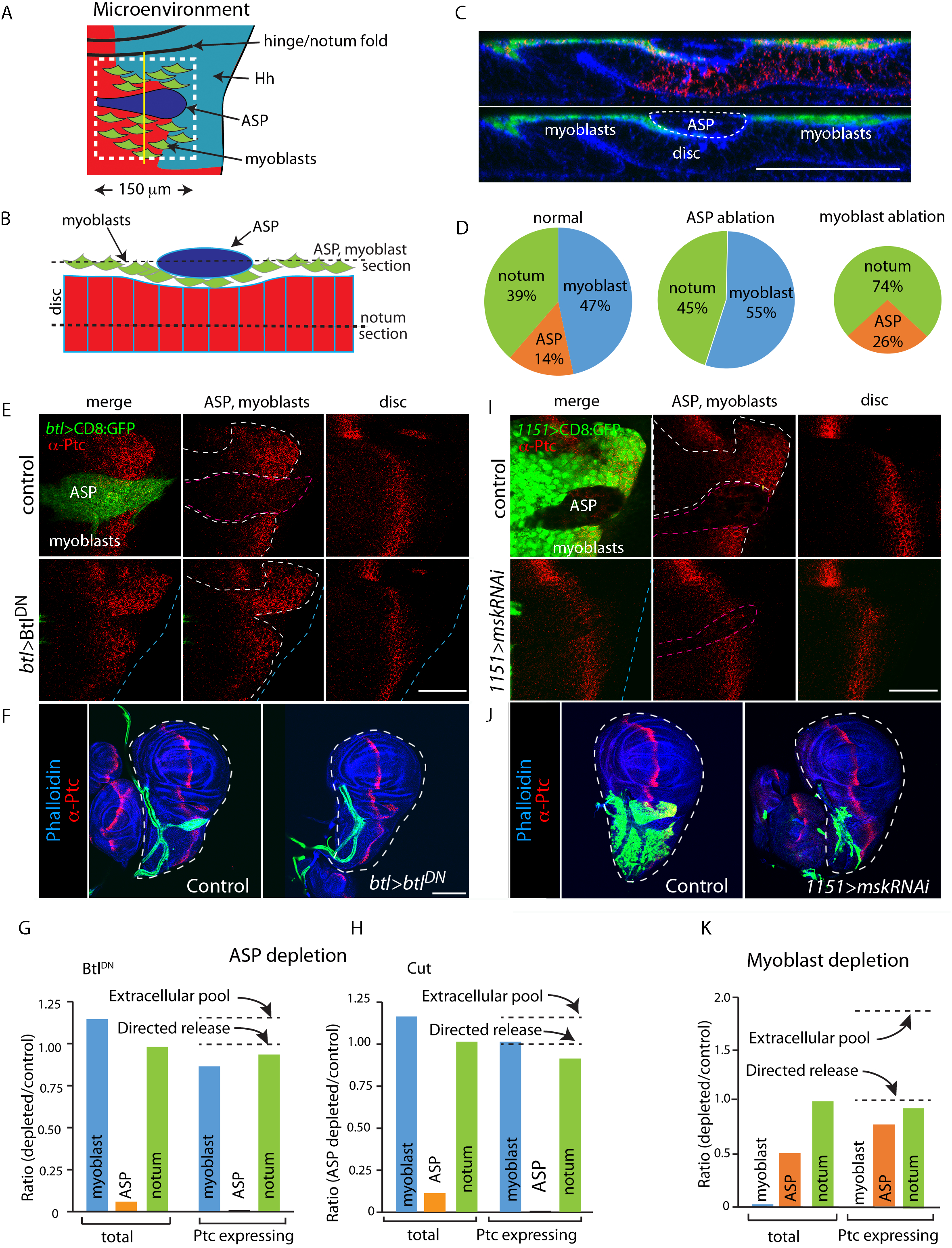
Hh distributions in the ASP, myoblast, and the notum primordium are not inter-dependent. (A) Schematic showing the microenvironment (white dashed line) in the notum primordium with myoblasts (green), ASP (blue), Hh expressing notum cells (turquois), and notum anterior compartment (red). (B) Schematic showing cross-section of the microenvironment at the yellow line in (A). (C) Confocal image of the cross section shown in (B); myoblasts (green), phalloidin staining (blue), α-Ptc staining (red). Scale bar: 50μm. (D) Pie graphs showing the percentage of microenvironment that expresses Ptc in the notum, myoblasts, and ASP under normal conditions and in conditions of ASP and myoblast ablation. (E) α-Ptc staining (red) of the microenvironment for control genotype (WT, ASP marked by CD8:GFP (green) driven by *btl-Gal4*; outlined by red dashed line in middle panel) and ASP ablation genotype (*btl-Gal4>*Btl^DN^); white dashed lines surround myoblasts, blue dashed lines indicate the notum primordium, yellow dashed lines mark the compartment boundary, blue arrows mark the extent of myoblast Ptc expression, green arrows mark extent of notum Ptc expression. Left column: ASP, myoblast section in (B), α-Ptc staining (red) and GFP (green); middle column: α-Ptc staining in the ASP, myoblast section indicated in (B); right column: α-Ptc staining in the notum section indicated in (B). Scale bar: 50μm. (F) Wing discs stained with α-Ptc antibody (red) and phalloidin (blue) for control genotype (WT) and ASP ablation genotype (*btl-Gal4>*Btl^DN^); trachea and ASP marked by CD8:GFP (green) driven by *btl-Gal4*. Scale bar: 100μm. (G) Bar graphs quantifying total area and Ptc-expressing areas of myoblasts, ASP, notum (total) in control (WT) and ASP-ablation genotype (*btl-Gal4>*Btl^DN^); n = 5 and 4 (control and ASP ablation, respectively). Dashed lines indicate predicted changes in Ptc expressing area under extracellular pool model of dispersion or directed release model of dispersion. (H) Similar to (G) but with the ASP depleted by the expression of Cut in the ASP (*btl-Gal4>cut*); n = 5 and 4 (control and ASP ablation, respectively). (I) Similar to (E) but with myoblast ablation; myoblasts marked CD8:GFP (green), ablated by knockdown of *msk* (*1151-Gal4>mskRNAi*). Orange arrows mark extent of Ptc expression in the ASP. Scale bar: 50μm. (J) Similar to (F) but with myoblast ablation. (K) Similar to (G) and (H), but with myoblast ablation; n = 5 for each genotype. Genotypes: (C) *1151-Gal4/+*; *UAS-CD8:GFP/+*; (E-G) control (*btl-Gal4 UAS-CD8:GFP/+*); *btl*>Btl^DN^ (*btl-Gal4 UAS-CD8:GFP/UAS-Btl*^*DN*^); (H) *btl*>Cut (*btl-Gal4 UAS-CD8:GFP/UAS-Cut; Gal80*^*ts*^*/+*); (I-K) control (*1151-Gal4/+*; *UAS-CD8:GFP/+*; *1151*>*mskRNAi* (*1151-Gal4/+*; *UAS-CD8:GFP/UAS-mskRNAi*).

We investigated the behavior of Hh that the three cell populations in this microenvironment share, testing if signaling in one tissue is influenced by the amount of Hh the others take up. The experiment distinguishes whether regulated Hh export creates a common pool of signaling protein that is shared among target cells, or if export is independently directed to target cells. If uptake is from a common pool, reducing the number of target cells is expected to increase uptake and signaling in the target cells that remain. Experimental addition of pools of extracellular and diffusible morphogens show that target cells are capable of responding to increases in morphogen available for uptake. Examples include both overexpression of a nonlipidated form of Hh (HhN) in the wing disc and addition of FGF-soaked beads in the chick limb bud increases signaling in the target cells (Callejo et al., 2006; Cohn et al., 1995).

We ablated the ASP and myoblasts genetically, and monitored Ptc expression as a readout of Hh signaling in the remaining tissues. The ASP does not develop when tracheal cells overexpress Btl^DN^ (Du et al., 2018; Sato and Kornberg, 2002), a dominant negative mutant FGFR protein (Reichman-Fried and Shilo, 1995), but the presence of Btl^DN^ in the tracheal cells has no apparent effect on the growth and morphogenesis of the disc or myoblasts (Fig. 7Supp A,B,E). In the absence of an ASP and therefore of the Hh target cells in the ASP, the amounts of Ptc in the disc and myoblasts were indistinguishable from controls (Fig. 7E-G, Fig. 7Supp G). We also ablated the ASP by ectopically over-expressing Cut, a transcription factor that negatively regulates FGF signaling (Du et al., 2018; Pitsouli and Perrimon, 2013). Ablation of the ASP by Cut expression similarly has no effect on the growth of the disc or myoblasts (Fig. 7Supp A,B), and it does not alter amounts of Ptc in the disc or myoblasts (Fig. 7H, Fig 7Supp G). To ablate myoblasts, we ectopically expressed *moleskin* (*msk*) RNAi. Msk is a nuclear importer of the FGF transcriptional activator ERK (Vishal et al., 2017). *mskRNAi* expression in the myoblasts reduced the number of myoblasts (Fig. 7D,I-K, Fig 7Supp D), and decreased the size of the ASP. This ASP phenotype is consistent with our previous findings that ASP growth and morphogenesis is dependent on Notch signaling from the myoblasts (Huang and Kornberg, 2015). Whereas the total amount of Ptc in the reduced population of myoblasts decreased under conditions of *msk* expression, disc growth and morphogenesis were unchanged, and Ptc expression in the disc was indistinguishable from controls (Fig. 7K, Fig. 7Supp F).

These results show that in the disc and myoblasts, Hh signaling is not dependent on Hh uptake by the ASP, and despite the fact that the number of Hh-responding myoblasts in the microenvironment was 20% greater than the number of Hh-responding disc cells at the stage these experiments were conducted, Hh signaling in the disc was not dependent on Hh uptake by myoblasts. We conclude that the number of Hh target cells in the ASP and myoblasts does not change the delivery of Hh to wing disc cells, and therefore that the spatial patterns of Hh signaling and Hh transport in the disc form independently of the presence or absence of other target cells.

## Discussion

This work shows that delivery of morphogen signaling proteins to target cells is controlled independently of both production amounts and target field size. In other systems, protein secretion has been characterized as either constitutive such that synthesis and discharge are concurrent and linked, or regulated such that synthesized proteins are made and stored until a stimulus instigates expulsion (Kelly, 1985). Antibody-producing lymphocytes are examples of constitutive secretory cells (Holodick et al., 2010). Examples of cells that regulate secretion include endocrine pancreatic cells that secrete hormones into the circulatory system and neurons that pass signals to target cells at synapses. The findings described here demonstrate that the distribution of Hh, Dpp, and Wg to target tissues has key attributes of regulated, targeted secretion.

Our previous work identified and highlighted many features of cytoneme-mediated signaling that are analogous to neuronal signaling. These include signal exchange at synapses that connect cells at distances of <40 nm (Roy et al., 2014), synaptic localization of proteins such as the Voltage-gated calcium channel and Synaptotagmin, essential roles for the glutamate receptor and glutamate transporter, and trans-synaptic stimulation of calcium transients (Huang et al., 2019). Cytoneme synapses are glutamatergic and calcium-dependent release of glutamate is essential for signaling by target cells. We consider the new properties of cytoneme-mediated Hh, Dpp, and Wg signaling that this work discovered in the context of this neuronal model. Single neurons produce and are sources of particular neurotransmitters. Neuronal signaling is spatially specific because neurotransmitters transfer to target cells at synapses that directly link neurons to pre-selected targets. And neuronal signaling is quantitatively precise because neurotransmitter release is controlled by regulated voltage-dependent discharge and programmed elimination after secretion. The distribution of neurotransmitter between signaling cells is asymmetric: most neurotransmitter is present inside neurons and is localized to presynaptic terminals poised for release. For example, the concentration of the neurotransmitter glutamate in vertebrate brain neurosecretory vesicles is 100-300 mM, but the fraction released in response to an action potential is small and extracellular concentrations are nanomolar (Burger et al., 1989; Chiu and Jahr, 2017; Riveros et al., 1986).

Our findings indicate that signaling proteins are also not released in amounts that are proportional to production. In the wing disc, Hh and Wg are made in greater quantities than are present in the cells they target (∼20x and 5x, respectively), the amounts of Hh, Dpp and Wg in target cells are insensitive to changes in the amounts of signaling protein that are generated by producing cells. We interpret this finding to indicate that these signaling proteins are not released constitutively, and from the perspective of the neuronal model, that their release is likely gated. Morphogen signaling protein production is cell-type specific, although it is generally spatially confined to contiguous groups of cells (“signaling centers”) that collectively express a signaling protein under temporal control. Signaling proteins distribute to nearby target cells as concentration gradients that decline with increasing distance from signaling center source cells. Although the capacity of cells to take up and respond to signaling protein is uniform over the target field in which the gradient forms, feedback regulation may influence the shape of a gradient. Hh signaling, for example, enhances expression of Ptc, a receptor protein that binds and sequesters Hh, and by reducing its spread, Ptc shapes the contour and extent of the Hh gradient. Importantly, we found that Ptc expression and the Hh gradient are insensitive to changes to the amount of Hh production (Fig. 1D,E). This implicates release as the step that controls the amount of Hh that distributes to target fields.

We consider several possible steps in the producing cell that might be regulated and rate-limiting. Post-translational modifications of Hh, Wg, and Dpp include proteolysis and lipidation, and are possibilities (Farzan et al., 2008; Künnapuu et al., 2009; Parchure et al., 2018). In addition, Hh and Wg traffic intracellularly between the apical and basolateral compartments and are sequestered in intracellular vesicles prior to release (Callejo et al., 2011; Gradilla et al., 2018; Yamazaki et al., 2016). Any of these and other steps that prepare signaling proteins for engagement with their respective receptors on a target cell might in principle be rate-limiting. Such mechanisms place the gating of delivery at a step at or prior to release, but do not distinguish if release is constitutive but rate-limited, or controlled. The context of release – whether it is at a site of a synaptic cell-cell contact or into extracellular fluid – is relevant to this question. The setting of Hh signaling in the wing disc that involves producing cells in the disc and recipient signaling cells in the ASP, myoblasts, and disc (Fig. 7) provided a way to answer it.

In the late third instar, the ASP extends posteriorly across the disc basal surface, from the far anterior to the A/P compartment border in the region just dorsal to the wing blade primordium. Myoblasts cover most of the disc A compartment in this region and extend over a portion of the P compartment as well. The myoblasts adhere to the disc and the ASP cells are juxtaposed directly to the disc in some places and in others are juxtaposed to myoblasts that adhere to the disc. All ASP, myoblast, and disc cells that are within approximately 62 μm of the Hh-expressing, P compartment disc cells activate Hh signaling (Hatori, R. and Kornberg, 2020). Thus, despite the differences in cell cycle, shape, constitution, and fate between the cells in this microenvironment, the primary determinant of Hh signaling is distance from source cells. If Hh were released into the extracellular space within this microenvironment and if the ASP, myoblast and disc cells in this microenvironment shared available Hh, we would expect that the distance over which Hh spreads as well as the extent of Hh signaling would depend on the number of recipient ASP, myoblast, and disc cells. It does not. Genetic ablation conditions that eliminate the myoblasts and reduce the number of cells in the target field by more than one-half does not increase the area in which disc cells activate Hh signaling (Fig. 7I-K). This indicates that the amount of Hh available to remaining cells does not increase, that Hh release is not rate-limiting, and that Hh is not released to a common pool that supplies Hh to cells in the microenvironment. Hh uptake in the microenvironment is determined by distance and not by production amounts or cell type.

If morphogen release is controlled and not constitutive, if release is not dependent on the number of target cells, if cells in the target field are equally capable of uptake and signaling, and if uptake is not from a common pool, how do spatial concentration gradients form? Contact-dependent, cytoneme-mediated spread is a likely mechanism because recipient cells that are closer to source cells make more cytoneme contacts (Du et al., 2018). Short cytonemes that connect nearby cells are more frequent than are long cytonemes that connect more distant cells, the consequence being that the number of synaptic contacts decreases with increasing distance. For FGF signaling in the ASP, positive feedback that increases the number of cytonemes of signaling cells and negative feedback that decreases the number of cytonemes in cells that do not signal may contribute to a spatial gradient of cytoneme number (Du et al., 2018). We suggest that similar feedback systems may sculpt the cytoneme gradients that disperse Hh and Dpp.

## Acknowledgements

We thank: H. Bellen and M. Gibson for antibodies and the Bloomington Stock Center and Vienna Drosophila Resource Center for fly stocks. This work was funded by NIH T32HL007185 to R.H and R35GM122548 to T.B.K.

## Materials and Methods

### Fly Genetics

#### Mutant lines

*hh*^*ac*^ (Lee et al., 1992), *dpp*^*H46*^ (Irish and Gelbart, 1987), *wg*^*RF*^ (Pérez-Garijo et al., 2009)

#### Transgenic lines

40k Hh BAC (Chen et al., 2017), 40k Hh:GFP BAC (Chen et al., 2017), 100k Hh BAC (Chen et al., 2017), Dpp BAC (this study), *Dpp:Cherry* crispr (Fereres et al., 2019), *wg-gal4* (Giráldez et al., 2002), *1151-gal4* (Roy and VijayRaghavan, 1997), *UAS-Wg:GFP* (Pfeiffer et al., 2002), *UAS-mcd8:GFP* (Roy et al., 2011), *btl-LHG* (Roy et al., 2014), *lexO-cherry:CAAX* (K. Basler), *UAS-Cut, UAS-btl*^*DN*^ (Reichman-Fried and Shilo, 1995), *UAS-mskRNAi* (Bloomington #27572)

#### Dpp BAC

BAC clone CH321-23O18 (CHORI) was inserted into cytological location 65B2 (Venken et al., 2009).

Flies were cultured in standard cornmeal and agar medium at 25°C and all crosses were done at 25°C, except expression of *Cut*. To express *Cut* in the ASP, *btl-Gal4*/*UAS-Cut*; *Gal80*^*ts*^/+ was incubated at 18°C until early L3 and transferred to 29°C until late L3.

### qPCR analysis of *hh* gene expression

Wing discs were dissected in PBS, RNA was extracted using RNeasy micro kit (Qiagen), and cDNA was synthesized using the High Capacity RNA-to-cDNA Kit (Applied Biosystems). qPCR was performed with SensiFast Sybr green (BIOLINE). For each genotype, 3-4 replicates of 5 wing discs were analyzed. Actin was the internal control and fold differences in relative mRNA levels between genotypes were calculated as 2^-ΔΔCt^.

### Immunohistochemistry, fluorescent imaging, and image analysis

Wing discs together with Tr2 trachea were dissected in PBS, fixed in 4% formaldehyde in PBS, and after washing in PBS-TritonX-100 (0.3%), samples were blocked in Roche Blocking Solution. Antibodies: mouse α-GFP (Roche), rabbit α-RFP (Rockland), mouse α-Ptc (DSHB, Apa1), mouse α-En (DSHB, 4D9), Rabbit α-Hh (from P. Ingham), α-pMAD (Abcam), α-Wg (DSHB), α-Sense (H. Bellen), α-Dpp prodomain (M.C. Gibson), Alexa633 conjugated-Phalloidin (Invitrogen), secondary antibodies (Invitrogen). Samples were mounted in Vectashield (Vector labs).

To observe cytonemes, unfixed preparations were observed using the hanging drop method (Huang and Kornberg, 2016). Images were acquired using the FV3000 Olympus Confocal microscope with GaAsP PMT detectors. Images were analyzed and processed with ImageJ and Photoshop. The domains of α-Ptc, α-Sense, α-En, and α-pMAD staining were measured in ImageJ from single optical sections at the basolateral part of the wing disc.

#### Statistics

In all figures, error bars indicate standard deviation (SD) and statistical significance was calculated by student’s t-test.

### Areas of Ptc expression

For Figure 6 (G,H,K), the dotted lines for the predicted extracellular pool was calculated assuming that total area of Ptc expression is constant and increase in remaining tissues compensates for the absence of ablated tissue. The ratio for directed release was set at 1.00 based on the assumption that Ptc expressing area for the remaining tissues would not change under conditions of tissue ablation.

#### ASP ablation

Calculated ratio of depleted/control for extracellular pool = 1.00/(% of Ptc expressing area in notum (0.39) + myoblast (0.47) before ablation) = 1.16

#### Myoblast ablation

Calculated ratio of depleted/control for extracellular pool = 1.00/(% of Ptc expressing area in notum (0.39) + ASP (0.14)) = 1.89

### Intensity measurements of proteins

Average intensity quantification for projections of α-Hh, α-GFP, α-Cherry, and α-Wg staining was calculated for segments of optical sections spanning 9 μm from the most apical side of the wing pouch cells. This segment was were chosen because the basal sides of the wing discs is folded, making accurate comparisons between samples problematic. Background measurements taken in equivalent areas distant from staining regions and were subtracted. Intensities in the Wg-producing and adjacent receiving cells were measured in the indicated 30 μm x 90 μm area, with the producing area defined by a 30 μm x 10 μm rectangle and the receiving area defined by two 30 μm x 40 μm rectangles.

### Quantification of cytoneme density

Maximum intensity projection image of the whole volume of the ASP was used to count the number of cytonemes in the bulb. To calculate the density of cytoneme per μm, the number of cytonemes was divided by the perimeter of the bulb of the ASP.

